# Adolescent development of anxiety-related behavior and shifts in behavioral responsiveness to estradiol in female mice

**DOI:** 10.1101/2024.09.02.610911

**Authors:** Kalynn M. Schulz, Marcia C. Chavez, Zoey Forrester-Fronstin

## Abstract

Early pubertal onset during adolescence is consistently linked with increased risk of anxiety and depression in girls. Although estradiol tends to have anxiolytic effects in adulthood, whether sensitivity to estradiol’s anxiolytic actions increases during adolescence is not clear. Using a rodent model, the current study tested the hypothesis that a shift in sensitivity to the anxiolytic effects of estradiol occurs during adolescence. To test this hypothesis, prepubertal and adult C57BL/6 female mice were ovariectomized, implanted with vehicle- or estradiol-filled silastic capsules, and behavioral tested one week later in the open field and elevated zero maze. Our hypothesis predicted that estradiol would decrease anxiety-related behavior to a greater extent in adults than in adolescent females, however, our results did not support this hypothesis. In the open field, estradiol implants significantly decreased anxiety-like behavior in adolescent females (relative to vehicle) and had little to no effect on the behavior of adults. These data suggest that adolescence is associated with a downward shift in sensitivity to the anxiolytic effects of estradiol on behavior in the open field. In contrast, although estradiol treatment did not influence anxiety-like responses in the elevated zero maze in early adolescent or adult females, adolescent females displayed significantly higher levels of anxiety-like behavior than adults. These findings demonstrate that substantial changes in anxiety-related behavior occur during adolescence, including a context-dependent shift in behavioral responsiveness to estradiol.

## 1. Introduction

The dramatic cognitive, social, and emotional changes that occur during adolescence bring both developmental opportunities and risks. Indeed, adolescence is typically when mental health disorders first emerge (de Lijster et al., 2017; Solmi et al., 2022), and females are nearly twice as likely as males to experience mental health difficulties during adolescence (Altemus et al., 2014; Kessler et al., 2012). A complex interplay of psychosocial and biological processes influences the likelihood of a healthy transition through adolescence. The pubertal onset of gonadal secretions of steroid hormones is a salient biological process that occurs during adolescence, and studies employing rodent models have advanced scientific knowledge of the organizational impact pubertal hormones exert on the developing adolescent brain (Brown and Spencer, 2013; Juraska and Willing, 2016; Piekarski et al., 2017; Schulz et al., 2009; Schulz and Forrester-Fronstin, 2018). Yet, our understanding of how pubertal hormones influence emotional behavioral development during adolescence remains limited.

To date, only a few animal studies have precisely tracked changes in anxiety-related behavior across adolescent development, and even fewer have included female subjects in their research designs. The results of these studies indicate that anxiety-like behavior *decreases* between early adolescence and adulthood in rats (Bishnoi et al., 2021; Graf et al., 2023; Lynn and Brown, 2010, 2009), and mice (Hefner and Holmes, 2007; Reiber et al., 2022). In adulthood, anxiety-related behavior is influenced by ovarian secretions, and many studies indicate that estradiol has anxiolytic effects on behavior. For example, clinical and preclinical studies report anxiolytic effects of estradiol during the high-estrogen phases of the menstrual cycle in humans (Farage et al., 2008; Ivey and Bardwick, 1968), and estrous cycle in rodents (Marcondes et al., 2001; Pestana and Graham, 2024). In addition, adult ovariectomy and estradiol treatment decreases rodent anxiety-like behavior in some behavioral assessments (Kastenberger et al., 2012; Renczés et al., 2020; Tomihara et al., 2009). Although estradiol modulates anxiety-related behavior in adult females, whether ovarian secretions of estradiol contribute to adolescent decreases in anxiety-like behavior is currently unknown. Likewise, whether behavioral neural circuits are immediately responsive to estradiol at the onset of puberty or acquire responsiveness during adolescent development is also unknown. To distinguish between these possibilities, we tested the hypothesis that the anxiolytic effects of estradiol on behavior develops during adolescence in female mice. Utilizing the open field and elevated zero maze, our strategy was to compare the behavioral responses of ovariectomized adolescent and adult females implanted with either estradiol- or vehicle-containing silastic capsules. Our hypothesis predicted that estradiol decreases anxiety-like behavior to a greater extent in adult than in early adolescent females.

## 2. Methods

### 2.1 Subjects

Timed pregnant C57BL/6 mice were purchased from Envigo (Indianapolis, IN) and arrived at the University of Tennessee Knoxville animal facility on day 2 of gestation. Dams and pups were maintained on a 12:12 light/dark cycle throughout the duration of the study and housed in polycarbonate cages (11.5” x 7.5” x 5”, 63 in^2^). In rodents, environmental sources of phytoestrogens affect sexual development, particularly in females. For example, isoflavone, a type of phytoestrogen, advances pubertal onset, disrupts normative estrous cycling, and alters responsiveness to gonadal steroid hormones (Takashimasasaki et al., 2007). To control for exogenous sources of estrogens in the bedding, food, and water, pregnant females and their offspring were provided with *ad libitum* access to phytoestrogen-free food (2020X Teklad global soy protein-free), bedding (Teklad shredded aspen), and BPA-free bottles. Female offspring were weaned on postnatal day (PD) 21 and housed 2 per cage with a same-treatment littermate. All procedures were approved by and carried out in accordance with the University of Tennessee Institutional Animal Care and Use Committee.

### 2.2 Determination of Pubertal Onset

The timing of pubertal onset under phytoestrogen-free housing conditions was assessed using a separate group of 20 gonadally intact females (Figure 2). Beginning at PD 25, vaginal opening was monitored daily during the last 2 hours of light in their light/dark cycle. Vaginal opening was not observed until day 27, which informed the timeframe for prepubertal ovariectomy in the experimental groups.

### 2.3 Experimental Design and Timeline

See Figure 1 for a timeline of experimental procedures. Prepubertal (PD 23-25) and adult (PD 73-75) females were ovariectomized (OVX) and implanted with silastic capsules filled with either sesame oil alone (vehicle) or estradiol dissolved in sesame oil. One week following ovariectomy, adolescent and adult females were tested for anxiety-related behaviors in the Open Field and Elevated Zero Maze (EZM) at 29-31 and 79-81 days of age, respectively (Figure 1). The adolescent age range at behavioral testing corresponds with the peripubertal timeframe between vaginal opening and first vaginal estrus in C57BL/6J mice that occurs in early adolescence (Piekarski, Boivin, et al., 2017).

**Figure 1.**
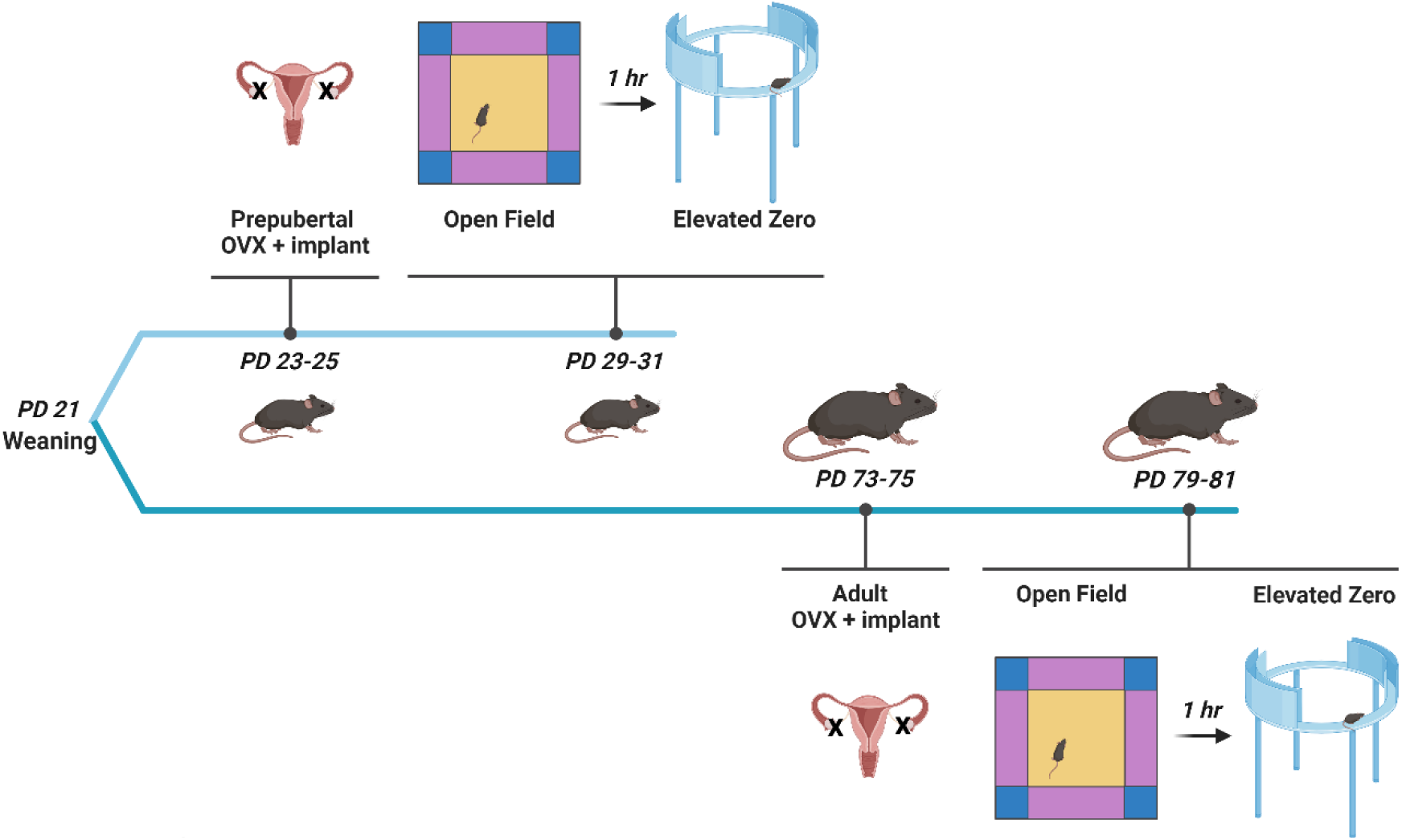
Timeline of experimental procedures. Females were ovariectomized and implanted with silastic capsules either prior to pubertal onset between 23-25 days of age, or in adulthood between 73-75 days of age. Half of all prepubertal and adult females received an estradiol-filled silastic capsule, and half received a vehicle-filled silastic capsule. One week following surgery, anxiety-related behavior was assessed in the Open Field and Elevated Plus maze. PD= postnatal day of age. Created with BioRender.com.

**Figure 2.**
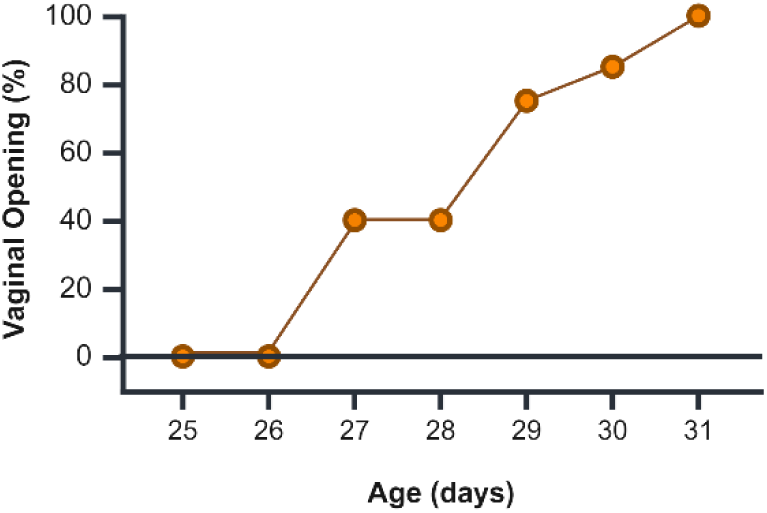
Timing of vaginal opening in 20 gonad-intact female mice (non-experimental). Vaginal opening was first observed on postnatal day 27 in 40% of females, and in 100% of females by postnatal day 31. These data informed the timing of prepubertal gonadectomy in experimental groups. Created with BioRender.com

### 2.4 Ovariectomy and silastic capsule implants

All surgical procedures were performed aseptically under isoflurane anesthesia. Pre-operatively, buprenorphine (0.03 mg/ml) and ketoprofen (0.1 mg/kg) were administered subcutaneously. Surgical preparations included shaving the dorsal hind flanks, sterilizing the incision sites with swabs of betadine and alcohol, and applying ophthalmic ointment to the eyes. The uterine horn and ovaries were located through a 3 mm incision in the skin and muscle wall, a hemostat was applied between the ovaries and uterine horn, and the ovaries were removed using a cautery instrument. The muscle wall incision was closed with silk braided non-absorbable suture (DT-719-1, DemeTECH) and the skin was closed with wound clips. Immediately following ovariectomy, females received a silastic capsule implant (Dow Corning Brand) containing either sesame oil alone (Spectrum, Hain Celestial Group LLC) or 17β-Estradiol (Eβ) dissolved in sesame oil (136 μg/mL, MP Biomedicals, LLC, 02101656.2). Silastic capsules were inserted subcutaneously between the shoulder blades through a 3 mm incision, and the incision was closed with wound clips (Reflex 7, Cellpoint Scientific, Inc., 203-1000). The capsule dimensions (inner diameter = 1.57mm, outer diameter = 3.17 mm, length = 22 mm) and concentration of Eβ (136 μg/mL) simulate levels of serum estradiol associated with proestrus in mice (Ström, et al., 2012). Following surgery, females received a subcutaneous injection of sterile saline (0.3 mL) and recovered from anesthesia in a clean cage placed on a heating pad. To minimize post-surgical discomfort, a subcutaneous injection of ketoprofen (0.1 mg/mL) was administered 24 hours later at 2.0 mg/kg.

### 2.5 Behavioral Testing Procedures

Behavioral testing occurred during the light phase of the light/dark cycle. On the day of testing, females were transported from the animal facility and were acclimated to the behavioral testing suite for at least one hour before testing. To minimize stress, the subjects experienced the same handler for both open field and elevated zero tests, and experimenters video monitored testing progress from an adjacent room. All behavior tests were video recorded and analyzed using Topscan behavioral analysis software (Clever Sys. Inc., Reston, VA). Simple Green Pro HD deodorizer (Huntington Harbour, CA) was used to clean the testing apparatus between subjects.

#### 2.5.1 Open Field Test

Indices of anxiety-related behavior were assessed using an open field arena. The open field arena was constructed of blue plastic (60cm x 60cm x 60cm). Mice were gently lowered into the arena perimeter at a standardized location and allowed to explore freely for five minutes. Specialized behavior recognition software (Topscan, Cleversys Inc.) allowed the arena to be visually split into a center zone (60% of the floor) with a surrounding perimeter (40% of the floor). The behavioral analysis software monitored the number of entries, duration, distance traveled, and velocity in the center and perimeter zones. Center investigation/sniffing from the perimeter zone was also recorded. Specifically, this measure required the animal’s hind feet and body (center point) to be in the perimeter when sniffing the center zone.

#### 2.5.2 Elevated Zero Maze

The circular EZM was constructed from blue plastic with a track width measuring 52cm and walls measuring 50 cm in height in the closed areas (Clever Sys, Reston, VA). Females were gently placed in the closed wall and allowed to explore for 5 minutes while being video recorded. The behavioral analysis software monitored open area entries, durations, and velocities in the open and closed wall areas of the zero maze. Sniffing behavior was also recorded when an animal sniffed the open area from the confines of the closed area (hind paws in the closed area).

### 2.6 Data Analysis

IBM SPSS version 29 was utilized for all statistical analyses. Three-factor mixed ANOVAs (Age x Implant x Zone) were conducted to evaluate the effects of Age and Implant across different zones of the Open Field or Elevated Zero Maze. Significant 3-way interactions were followed by simple interaction and simple main effects analyses. Two-factor between-subjects ANOVAs focused solely on the center zone (Open Field), or open area (Elevated Zero) were also conducted. All interactions were interpreted using simple comparisons of the estimated marginal means, 95% confidence intervals associated with mean differences, and estimates of effect size (partial eta squared, ηp^2^). Partial eta squared values were interpreted as small (ηp^2^, 0.01-0.059), moderate (ηp^2^, 0.06-0.139), or large (ηp^2^ 0.14 and greater) effects. The relationship between thigmotaxis and velocity was investigated using Pearson’s *r.* In all cases, statistical significance was considered *p* ≤ 0.05.

## 3. Results

### 3.1 Vaginal opening in gonad-intact females

Twenty gonadally intact females were observed for vaginal opening beginning at PD 25 (Figure 1). Eight females exhibited vaginal opening at 27 days of age (40%). By PD 29, vaginal opening was observed in 16 females (80%), and all females exhibited vaginal opening by postnatal day 31 (100%).

### 3.2 Open Field

#### 3.2.1 Percent Time in Zones of the Open Field

We first sought to verify the anxiogenic nature of the center zone by analyzing the percent time females spent in the center, midsections, and corner zones of the open field (Figure 3). The 3-factor mixed ANOVA revealed a significant main effect of Zone (*F(*2,43) = 1676.40, *p* < 0.001, ηp^2^ = 0.99) such that irrespective of Age or Implant, subjects spent significantly less time in the center zone than either the midsection (*Mean Difference* = −32.98%, CI 95% [−34.63% - 31.33%], *p* < 0.001) or corner zones (*Mean Difference* = −48.73%, CI 95% [−52.21% - −45.26%], *p* < 0.001), confirming the center zone was anxiogenic. There was also a significant interaction amongst Age, Implant, and Zone (*F*(2,43) = 3.4, *p* = 0.044, ηp^2^ = 0.14). To evaluate the 3-way interaction, the effects of Age and Implant were probed separately for each zone of the open field: the center (Figure 3A), midsection (Figure 3B), and corner (Figure 3C). In the center zone, a simple main effect of Age revealed that adults spent a greater proportion of total test time in the center zone than did adolescents (*F(*1,44) = 6.63, *p* = 0.013, *Mean Difference* = 2.86, CI 95% [0.62% - 5.12%], ηp^2^= 0.13). Although Implant did not influence the proportion of center time (*F(*1,44) = 0.13, *p* = 0.72, ηp^2^= 0.003), Age and Implant significantly interacted (*F(*1,44) = 4.17, *p* = 0.047, ηp^2^ = 0.09,) indicating that differences between adolescents and adults depend on treatment with vehicle or estradiol implants. Specifically, a simple effect of age (adults > adolescents) was present in vehicle-implanted females (*Mean Difference* = 5.14%, CI 95% [1.83% - 8.44%], *p* = 0.003, ηp^2^ = 0.18), but not estradiol-implanted females (*Mean Difference* = −0.59%, CI 95% [−2.44% - 3.63%], *p* = 0.70, ηp^2^ = 0.004). In contrast to the center zone, no main effects or interactions were found in either the midsection or corner zones (Figure 3B & 3C). (Midsections Age: *F(*1,44) = 0.23, *p* = 0.64, ηp^2^ = 0.0; Midsections Implant: *F(*1,44) = 0.91, *p* = 0.35, ηp^2^ = 0.02; Midsections Interaction: *F(*1,44) = 0.29, *p* = 0.59, ηp^2^ = 0.01; Corners Age: *F(*1,44) = 0.63, *p* = 0.43, ηp^2^ = 0.01; Corners Implant: *F(*1,44) = 0.71, *p* = 0.40, ηp^2^ = 0.02; Corners Interaction: *F(*1,44) = 0.26, *p* = 0.61, ηp^2^ = 0.01).

**Figure 3.**
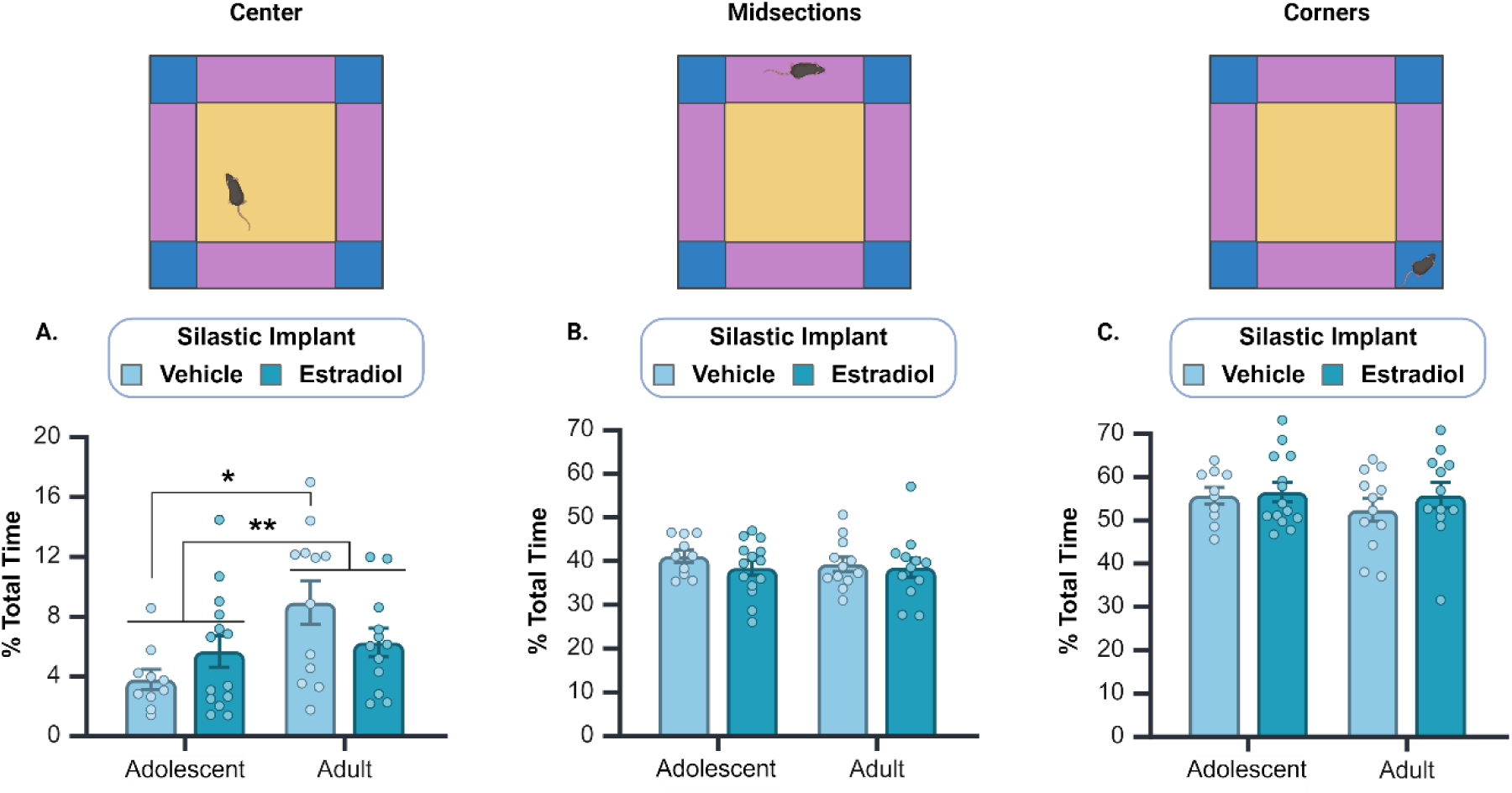
Percent of time during the 5-min Open Field Test spent in the center (A), midsections (B), and corners (C) of the open field. Given the anxiogenic nature of the center zone, all subjects spent the majority of test time in the midsections and corners of the open field. However, experimental effects were only present in the center zone (A). An interaction between Age and Implant was driven by a difference between vehicle-implanted adolescents and adults that was eliminated in estradiol-implanted adolescent and adult females. Circles represent individual values, and the bars represent mean +/- SEM. * denotes p < 0.05; ** denotes p < 0.01. Created with BioRender.com

#### 3.2.2 Open Field Center Entries and Bout Durations

Adults entered the center zone significantly more often than did adolescents (Figure 4A; *F(*1,44) = 6.14, *p* = 0.017, *Mean Difference* = 5.11, CI 95% [0.96 - 9.27], ηp^2^= 0.12). Implant did not affect the number of center entries (*F(*1,44) = 0.01, *p* = 0.94, ηp^2^ = 0.00), nor did Age and Implant interact to impact center entries (*F(*1,44) = 0.23, *p* = 0.63, ηp^2^ = 0.01). We also assessed the average bout duration of center entries and found a significant interaction between Age and Implant (Figure 4B; *F(*1,44) = 9.97, *p*= 0.003, ηp^2^ = 0.19). In adolescents, average bout durations were longer in estradiol-treated than in vehicle-treated females (*Mean Difference* = 0.71, CI 95% [0.21 - 1.21], *p* = 0.007, ηp^2^ = 0.12). In adults, average bout durations were nonsignificantly decreased in estradiol-treated females relative to vehicle-treated females, which likely contributed to the significant interaction (*Mean Difference* = −0.40, CI 95% [−0.89 - 0.10], *p* = 0.11, ηp^2^ = 0.06). No main effects of Age or Implant were present for this measure (Age: F(1,44) = 0.46, *p* = 0.50, ηp^2^ = 0.01; Implant: F(1,44) = 0.79, *p* = 0.38, ηp^2^ = 0.02).

**Figure 4.**
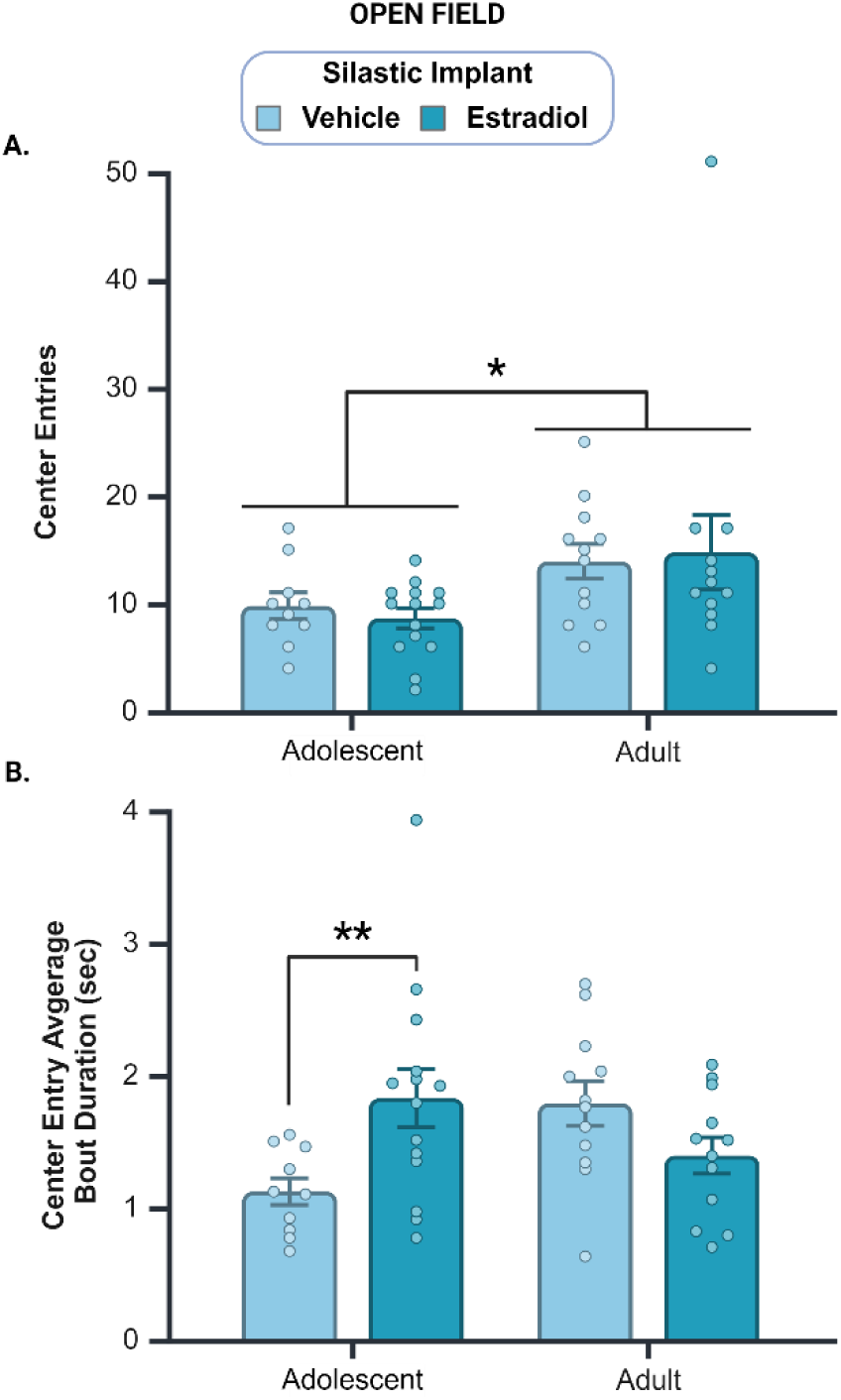
Center entry number and average bout duration in the Open Field Test. A. Adult females entered the open field center significantly more often than did adolescents. B. A significant interaction between Age and Implant was driven by differential effects of Implant in adolescents and adults. In adolescents, estradiol increased entry bout durations relative to vehicle-implanted subjects, whereas no significant differences were observed between estradiol- and vehicle-implanted adults. Circles represent individual values, and the bars represent mean +/- SEM. * denotes p < 0.05; ** denotes p < 0.01. Created with BioRender.com

#### 3.2.3 Velocity in the Open Field: relationship to anxiety-related behavior

We investigated whether velocity in the open field is associated with anxiety-like behavior (Figure 5A). Thigmotaxis is a well-validated marker of anxiety-like behavior in rodents (Scharf and Farji-Brener, 2024). Using perimeter duration as a proxy for thigmotaxis, Pearson product-moment correlation coefficients were computed to assess the relationship between perimeter duration and velocity in the center and perimeter zones. Perimeter duration and center velocity were moderately positively correlated (Figure 5A, *left panel; r*(48) = 0.60, *p* < 0.001, CI 95% [0.35 - 0.74]), indicating that subjects displaying higher levels of thigmotaxis also tended to travel through the center zone at higher velocities. In contrast, no correlation was found between perimeter duration and velocity in the perimeter (Figure 5A, *right panel*; *r*(48) = 0.21, *p* > 0.05, CI 95% [0.08 - 0.46]). The center zone-specific relationship between thigmotaxis and velocity supports the construct validity of utilizing center velocity as an anxiety indicator in the open field test.

**Figure 5.**
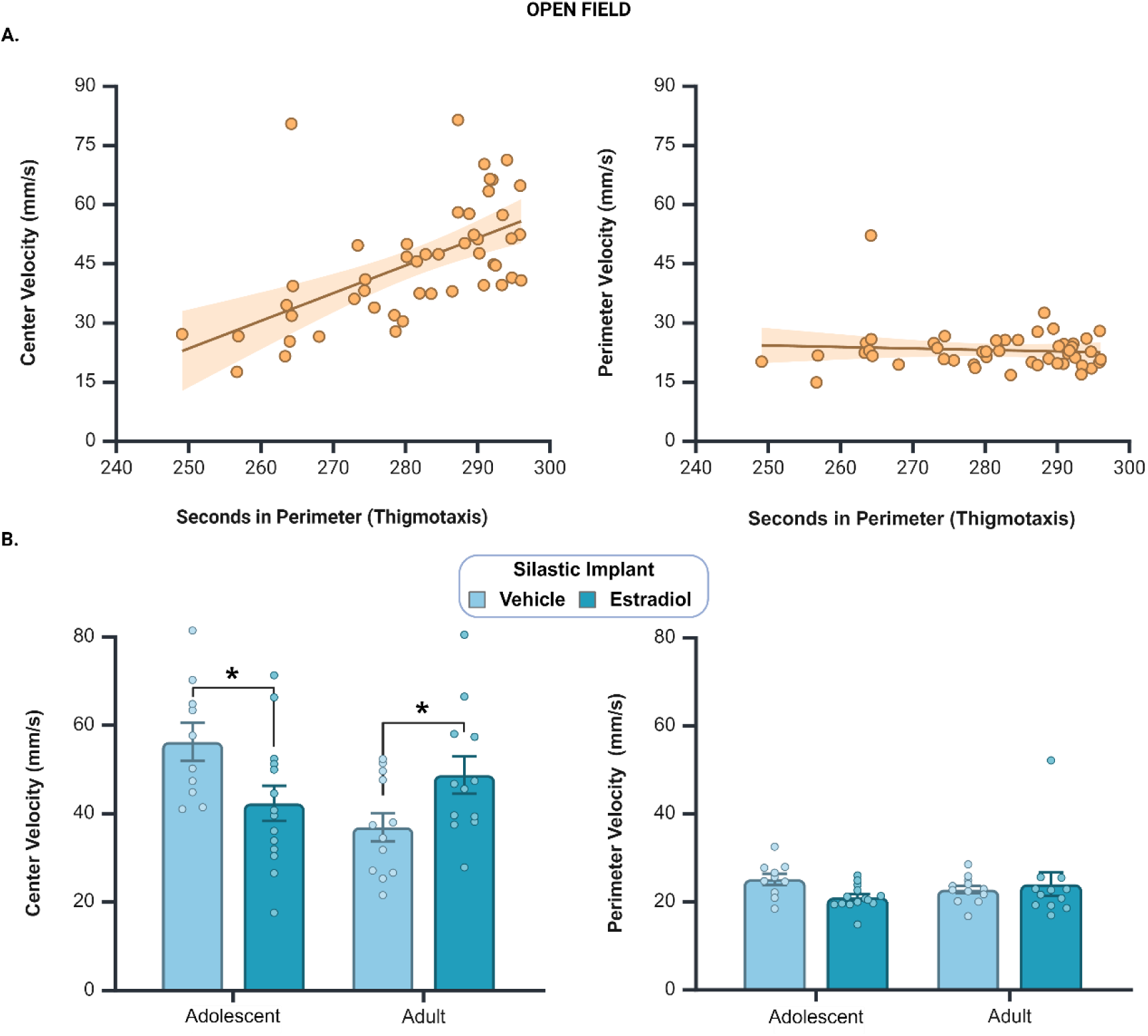
Velocity in the Open Field Test. A. To characterize the relationship between velocity and anxiety-related behavior in the Open Field, we tested whether center velocities correlate with a well-validated index of anxiety-related behavior, perimeter duration (thigmotaxis). Perimeter duration correlated positively center velocity, indicating that animals displaying higher levels of thigmotaxis exhibit higher center velocities (A, left panel). In contrast, no relationship was found between perimeter duration and perimeter velocity (A, right panel). These data suggest that velocity in the open field center is a meaningful measure of anxiety-like behavior in mice. Orange shading indicates 95% confidence intervals. B. A 3-factor Mixed ANOVA revealed a significant interaction between Age, Implant, and Zone (Center vs. Total Perimeter) on velocity. Simple interactions between Age and Implant were present in the center (B, left panel) but not in the perimeter Zone (B, right panel). In adolescents, estradiol significantly decreased center velocity relative to vehicle- implanted age-matched subjects. In adults, estradiol significantly increased velocity relative to vehicle-treated females. Circles represent individual values, and the bars represent mean +/- SEM. * denotes p < 0.05. Created with BioRender.com

#### 3.2.4 Velocity in the Open Field: Center Zone vs. Total Perimeter

Velocity in the center vs. the total perimeter was analyzed by 3-factor mixed ANOVA treating Age (adolescent vs. adult) and Implant (vehicle vs. estradiol) as between-subjects factors, and Zone (center vs. total perimeter) as the repeated measure. A main effect of Zone indicated that animals traveled at significantly higher velocities in the center relative to the perimeter Zone (Figure 5B, *left panel*; *F(*1,44) = 171.50, *p* < 0.001, *Mean Difference* = 22.80, CI 95% [19.30 - 26.31], ηp^2^ = 0.80). In addition to the main effect of Zone, a significant interaction between Age, Implant, and Zone (*F(*1,44) = 8.54, *p* = 0.005, ηp^2^ = 0.20) warranted investigation of velocity for the center and perimeter zones separately. In the center zone, Age and Implant interacted to impact velocity (Figure 5B, *left* panel; F(1,44) = 10.44, *p*= 0.002, ηp^2^ = 0.200). In adolescents, estradiol-implanted females displayed significantly *decreased* center velocities relative to vehicle-implanted females (*Mean Difference* = −13.92, *p* = 0.02, 95% CI [−25.324 - −2.513], ηp^2^ = 0.12). In contrast, adult estradiol-implanted females displayed significantly *increased* center velocities relative to vehicle-implanted adults (*Mean Difference* = 11.76, *p* = 0.04, 95% CI [0.52 - 23.01], ηp^2^ = 0.10). In the Perimeter zone, velocity was not impacted by Age or Implant, nor did these factors interact (Figure 5B, *right* panel; Age: *F(*1,44) = 0.05, *p* = 0.83, ηp2 = 0.001; Implant: *F(*1,44) = 0.86, *p* = 0.36, ηp^2^ = 0.02; Interaction: *F(*1,44) = 2.89, *p* = 0.10, ηp^2^ = 0.06).

### 3.3 Elevated Zero Maze

#### 3.3.1 Closed Wall Measures

Two measures were analyzed separately by two-factor ANOVA: (1) The duration of time spent sniffing open areas from the closed walls, and (2) the percentage of time within the closed walls spent sniffing the open areas. Both analyses revealed main effects of Age, indicating that adults spent longer durations sniffing the open area than did adolescents (Figure 6A; *F(*1,44) = 6.10, *p* = 0.02, *Mean Difference* = 11.84, CI 95% [2.18 - 21.50], ηp^2^ = 0.12), and also a greater percentage of time sniffing the open areas from the closed walls (Figure 6B; *F(*1,44) = 13.80, *p* < 0.001, *Mean Difference* = 7.0%, CI 95% [3.20% - 10.80%], ηp^2^ = 0.24). No effects of Implant, or interactions between Age and Implant were found for the duration sniffing the open areas or the percentage of closed wall time spent sniffing the open areas (Sniffing Open, Implant: *F(*1,44) = 0.18, *p* = 0.67, ηp^2^ = 0.004; Sniffing Open, Age x Implant: *F(*1,44) = 1.43, *p* = 0.24, ηp^2^ = 0.03; %Time Sniffing Open, Implant: *F(*1,44) = 0.55, *p* = 0.46, ηp^2^ = 0.01; %Time Sniffing Open, Age x Implant: *F(*1,44) = 0.71, *p* = 0.41, ηp^2^ = 0.02).

**Figure 6.**
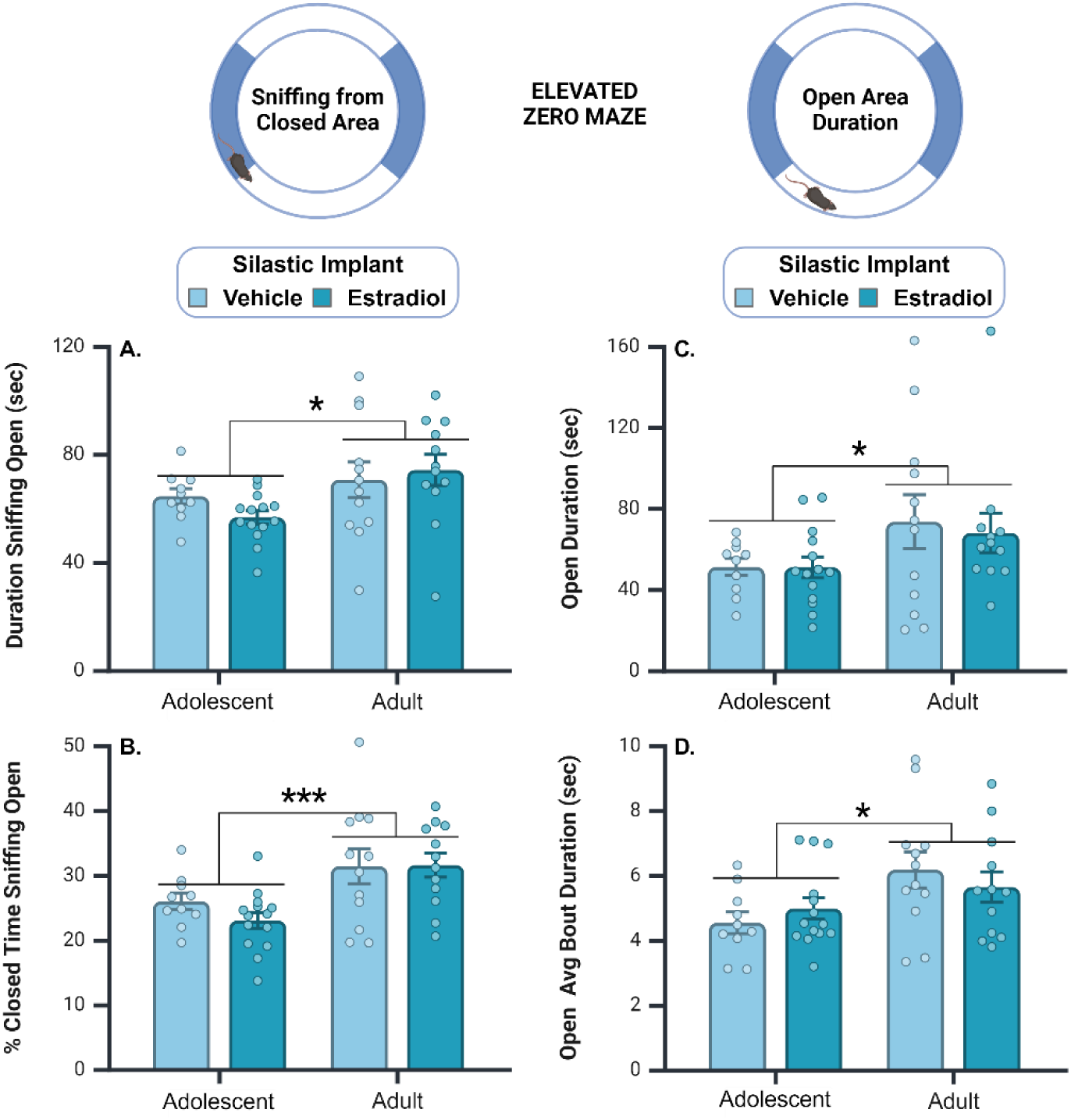
Open area investigation from the closed area and time spent in the open area of the Elevated Zero Maze. While in the closed wall area, main effects of Age indicated that adults investigated the open areas for longer durations (A) and spent a greater proportion of closed area time sniffing the open areas than did adolescent females (B). Duration in the open area (C) and open entry average bout durations (D) were also significantly greater in adult than in adolescent females. Circles represent individual values, and the bars represent mean +/- SEM. * denotes p < 0.05; *** denotes p < 0.001. Created with BioRender.com

#### 3.3.2 Open Area Measures

The total duration of open area entries and the average entry bout durations were analyzed separately by two-factor ANOVA. Adults spent overall longer durations in the open areas than did adolescents (Figure 6C; *F(*1,44) = 4.73, *p* = 0.035, *Mean Difference* = 19.65, CI 95% [1.45 - 38.00], ηp^2^ = 0.10). Similarly, adults displayed longer entry average bout durations than adolescents (Figure 6D; *F(*1,44) = 6.85, *p* = 0.012, *Mean Difference* = 1.15, CI 95% [0.26 - 2.03], ηp^2^ = 0.14. No effects of Implant, or interactions between Age and Implant were found for either measure (Open Duration, Implant: *F(*1,44) = 0.10, *p* = 0.76, ηp^2^ = 0.002; Open Duration, Age x Implant,: *F(*1,44) = 0.09, *p* = 0.77, ηp^2^ = 0.002; Open Average Bout Duration, Implant: *F(*1,44) = 0.01, *p* = 0.92, ηp^2^ = 0.00; Open Average Bout Duration, Age x Implant: *F(*1,44) = 1.24, *p* = 0.27, ηp^2^ = 0.03).

#### 3.2.3 Velocity in the Open Area and Closed Walls

We investigated whether velocity in elevated zero is associated with anxiety-related behavior. Using closed wall duration as a proxy for thigmotaxis, Pearson product-moment correlation coefficients were computed to assess the relationship between closed wall duration and velocity in the open and closed walls. Closed wall duration correlated negatively with velocity in both locations. Specifically, closed wall duration was weakly negatively correlated with velocity in the open area (*r*(48) = −0.32, *p* = 0.03 CI 95% −0.55 - −0.04]), and strongly negatively correlated with velocity in the closed walls (*r*(48) = −0.82, *p* < 0.001, CI 95% [−0.90 - −0.70]). These data suggest that individuals displaying higher levels of thigmotaxis behavior tend to move at lower velocities in both the open and closed areas of the elevated zero maze. Because maze location failed to influence the direction of the relationship between thigmotaxis and velocity, we concluded that velocity lacks construct validity as an anxiety indicator in the elevated zero maze, and no further analyses were conducted.

## 4. Discussion

The current study tested the hypothesis that a shift in sensitivity to the anxiolytic effects of estradiol occurs during adolescence in female mice. To test this hypothesis, prepubertal and adult females were gonadectomized, implanted with vehicle- or estradiol-containing silastic capsules, and behavioral tested one week later in early adolescence or young adulthood. Our hypothesis predicted that estradiol would decrease anxiety-related behavior to a greater extent in adults than in adolescent females, however, our results did not support this hypothesis. In the open field, estradiol implants significantly decreased anxiety-like behavior in adolescent females (relative to vehicle) and largely failed to influence the behavior of adults. These data suggest that adolescence is associated with a downward shift in sensitivity to the anxiolytic effects of estradiol on behavior in the open field. In contrast, although estradiol treatment did not influence anxiety-like responses in the elevated zero maze in early adolescent or adult females, adolescent females displayed significantly higher levels of anxiety-like behavior than adults. Taken together, our findings demonstrate that remarkable changes in anxiety-related behavior occur during adolescence, including an assay-dependent shift in behavioral responsiveness to estradiol.

To our knowledge, this is the first study to compare the effects of estradiol on anxiety-like behavior in early adolescence and adulthood. Although estradiol did not impact anxiety-like behavior in the elevated zero, the age-related decline observed is consistent with previous studies utilizing gonad-intact rodents (Bishnoi et al., 2021; Graf et al., 2023; Hefner and Holmes, 2007; Lynn and Brown, 2010; Reiber et al., 2022). For example, a comparison of male and female rat anxiety-like responses at early, mid, and late adolescence revealed stepwise behavioral decreases in both the open field and elevated plus maze (Lynn and Brown, 2010). A similar developmental decline has been reported in male C57BL/6J mice (Hefner and Holmes, 2007; Moore et al., 2011), but not in male or female CD2 mice (Macrí et al., 2002). It is unclear whether the different developmental patterns reported between studies in mice reflect strain-specific developmental trajectories or differences in the specific methods employed in each study. Notably, whether testing occurred during the active/light phase or inactive/dark phase of the light/dark cycle differed between these studies. Our findings in C57BL/6J females align with previous studies in C57BL/6J males that also tested subjects during the light phase of the light/dark cycle (Hefner and Holmes, 2007; Moore et al., 2011). Given that testing in the dark or light phase of the cycle influences the display of anxiety-like behavior in rats (Andrade, 2003; Huynh et al., 2011), additional studies are needed to investigate potential circadian influences on anxiety-like behavior in adolescent and adult mice.

Both the open field and elevated zero maze test take advantage of an animal’s thigmotaxis behavior, which is the natural tendency for animals to stay in close proximity to walls or vertical surfaces when exploring new environments (Scharf and Farji-Brener, 2024). The relationship between thigmotaxis behavior and anxiety has been pharmacologically validated in studies demonstrating that anxiolytic drugs decrease thigmotaxis behavior and increase time spent in the anxiogenic open areas of novel environments (Kulkarni et al., 2007; Macrí et al., 2002; Pickles and Hendrie, 2013; Scharf and Farji-Brener, 2024). Less well understood, however, is whether other aspects of locomotor behavior such as velocity may also reflect an animal’s emotional state. As an initial investigation, we tested the relationship between thigmotaxis behavior and velocity in both the anxiogenic and non-anxiogenic zones of the open field and elevated zero maze. If velocity is associated with anxiety-like behavior, the relationship between thigmotaxis and velocity should differ between anxiogenic and non-anxiogenic zones of the testing apparatus. A zone-specific relationship was found in the open field test. Specifically, a positive correlation between perimeter duration (thigmotaxis) and velocity was found in the anxiogenic center zone, but not in the perimeter zone. These data suggest that velocity in the open field center and perimeter zones may reflect unique emotional states, and that increased center velocity may indicate an increased anxiety-like state. In contrast, zone-specific relationships between thigmotaxis duration and velocity were not observed in the elevated zero maze. Experimental tests are needed to pharmacologically validate or refute a relationship between velocity and anxiety in the open field or elevated zero maze.

Currently, our scientific understanding of gonadal hormone influences on adolescent cognitive and emotional behavioral development trails behind understanding of adolescent sociosexual behavioral development (Lin and Wilbrecht, 2022). For sexual behavior, steroid hormone treatments that readily activate mating behavior in adult male and female rodents, fail to elicit behavioral responses prior to adolescence (Meek et al., 1997; Olster and Blaustein, 1989; Romeo et al., 2002, 2001), suggesting that behavioral neural circuits underlying sexual behavior acquire steroid hormone responsiveness during adolescent development. Interestingly, for anxiety-like behavior, our results in the open field suggest that behavioral responsiveness to estradiol *decreases* during adolescence. Thus, adolescent shifts in behavioral responsiveness to steroid hormones may be in opposite directions for sexual (low to high) and anxiety-like behavior (high to low). It is currently unknown whether male rodents also exhibit a change in anxiety-like responsiveness to testosterone before and after adolescence.

Shifts in behavioral responsiveness from pre- to post-adolescence indicate that developmental processes occurring *specifically during adolescence* are necessary for adult-typical behavioral responses to steroid hormones. For sociosexual behaviors, this shift in responsiveness is governed by the timing of intersection between pubertal gonadal hormones and the developing adolescent brain. For example, in males deprived of gonadal hormones during adolescence via prepubertal gonadectomy, adult testosterone replacement fails to elicit adult-typical levels of mating behavior. In contrast, in males gonadectomized after puberty, mating behavior is fully restored following testosterone placement in adulthood (Schulz et al., 2009, 2004). These data suggest that pubertal hormones shape developing neural circuits during adolescence and program adult behavioral responses to testosterone in adulthood. Accumulating evidence in male rodents suggests that adult anxiety-related behavior is also organized by gonadal steroid hormones during adolescence (Brand and Slob, 1988; Brown et al., 2015; Primus and Kellogg, 1990). Unlike sexual behavior, anxiety-related behavior is modulated by, but not dependent upon, the presence of gonadal hormones in males and females (Bitran et al., 1993; Carrier et al., 2015; Domonkos et al., 2017, 2017; Fernandez-Guasti and Martinezmota, 2005; Kastenberger et al., 2012; Marcondes et al., 2001; Näslund et al., 2016; Pestana and Graham, 2024; Renczés et al., 2020; Tomihara et al., 2009). Therefore, the effects of pre- and post-pubertal gonadectomy on adult anxiety-related behavior can be assessed in the presence and absence of steroid hormone replacement in adulthood. Indeed, males gonadectomized before and after puberty display marked differences in adult anxiety-like behavior *in the absence of testosterone replacement* (Brown et al., 2015). Whether adult anxiety-like responses to steroid hormones vary depending on adolescent gonadal hormone exposure is currently unknown. Scientific models of steroid hormone influences on adolescent behavioral development will benefit from studies in *both sexes* investigating the effects of steroid hormones on anxiety-like behavior (1) before, during, and after adolescence, and (2) in in animals gonadectomized before and after puberty and tested in adulthood.

Girls who undergo puberty at younger ages than their peers are at increased risk for mental health conditions such as anxiety and depression (Graber, 2013; Graber et al., 2010; Kowalski et al., 2021; Zehr et al., 2007). Considering that the age of pubertal onset is decreasing world-wide (Bellis et al., 2006; Brix et al., 2019; Eckert-Lind et al., 2020; Parent et al., 2003, 2016), understanding the mechanisms by which earlier pubertal timing impacts mental health outcomes is imperative. Our findings in the open field test suggest that adolescent females are sensitive to the anxiolytic actions of estradiol on behavior, and more sensitive than adult females. Shifts in neural and behavioral responses to steroid hormones across puberty and adolescence (Romeo et al., 2002; Terasawa et al., 2013), and indicate that processes occurring during adolescence change neural responses to gonadal steroid hormones. These shifts may be steroid-independent and developmentally timed, as in the case with the decreased hypothalamic-pituitary-gonadal axis sensitivity to steroid negative feedback during puberty (Sisk and Foster, 2004). These shifts in responsiveness may also be the consequence of steroid-dependent development of behavioral neural circuits during puberty, as is the case for sociosexual behaviors (Romeo, 2005; Schulz et al., 2009, 2006). Importantly, although we report here an ontogenetic shift in anxiety-like responses to estradiol during adolescence, additional studies are needed to identify the specific processes occurring during adolescence that give rise to changes in responsiveness. Studies utilizing animal models of early pubertal timing in females hold promise for uncovering these biological processes (Boivin et al., 2017), and will increase knowledge of how interactions between biological, social and environmental factors may contribute to increased risk for anxiety and depression during adolescence.

## 5. Author Contributions

**Kalynn Schulz**: Conceptualization, methodology, formal analysis, resources, data curation, review and editing, visualization, supervision, and funding acquisition.

**Marcia Chavez:** Methodology, formal analysis, investigation, data curation, original draft, and review and editing.

**Zoey Forrester-Fronstin:** Methodology, investigation, data curation, review and editing, and project administration.

## 7. Acknowledgements

We thank Abby Turner, Mackenzie Hooker, and Harry Pepper for their teamwork and excellent care of our research animals.

